# Cerebrospinal fluid flow and clearance driven by lateral ventricle volume oscillations

**DOI:** 10.1101/2025.10.20.683529

**Authors:** Viktor Neumaier, Moritz Bonhoeffer, Melissa Thalhammer, Julia Schulz, Gabriel Hoffmann, Juliana Zimmermann, Felix Brandl, Matthias Brendel, Igor Yakushev, Josef Priller, Jan Kirschke, Christine Preibisch, Benedikt Zott, Christian Sorg

## Abstract

In the human brain, substance clearance is intimately connected to the cerebrospinal fluid (CSF) and its flow. CSF extends from the lateral ventricles (LVs) to the parenchyma’s perivascular spaces. Macroscopic undulating CSF flow is present during both wakefulness and sleep and can be experimentally induced. However, the mechanisms generating this flow and its contribution to brain clearance remain unclear. Using fMRI and PET across various conditions, we demonstrate that LV-volume oscillations drive undulating CSF flow in the ventricles and subarachnoid basal cisternae. LV oscillations are driven by cortical blood-volume changes induced by neuronal activity, heartbeat and respiration. LV oscillations’ amplitudes determine PET-tracer clearance from the LVs. Conclusively, induced by extra- and intracranial physiological drivers and mediated by cortical blood-volume changes, LV-volume oscillations drive macroscopic CSF flow and clearance.

## Main Text

Cerebrospinal fluid (CSF) and its flow are crucial for cerebral substance clearance (*1, 2*). Primarily secreted by the ventricular choroid plexus (*3*), CSF travels from the lateral ventricles (LVs) to the subarachnoid space surrounding the brain. From there, according to the model of the glymphatic system, CSF enters the brain parenchyma along the perivascular spaces around penetrating arterioles, mixes with the interstitial fluid, and exits the brain via several outflow pathways, eventually into the neck’s lymphatic and venous vessels (*4, 5*). Along this CSF-glymphatic pathway, substances are distributed (*6*) and cleared (*7, 8*). Critically, for substance distribution and clearance to be effective, it must be supported by directional solute movement with clear in- and outflow sides. Furthermore, directional solute clearance needs to be driven by a moving solvent, presumably by the movement of CSF propelled through distinct physiological drivers.

In the microscopic pathways of the parenchymal periarterial spaces, bulk CSF movement is driven by vessel wall displacement induced by extracranial cardiac and respiratory (*9*) as well as intracranial neuronal activity (*10*). CSF movement in the macroscopic pathways from the LVs to the periarterial spaces is less understood. Based on the Monro-Kellie doctrine, which states that a stable sum of CSF-, blood-, and interstitial volumes is required within the fixed cranium to maintain constant intracranial pressure (*11*), global models of macroscopic CSF flow suggest that total brain blood-volume changes lead to anti-correlated CSF volume changes, causing CSF movement (*12*–*14*) into and from the cranium. The drivers of this flow include, again, extracranial oscillatory activity of heartbeat and respiration (*15*–*18*) and intracranial blood-volume changes induced by neuronal and vascular activity (*12, 14, 19, 20*).

However, how the oscillatory activity of these drivers mechanistically generates macroscopic CSF flow remains unclear. Moreover, it is also unknown how undulating CSF flow can support directional substance movement and clearance. A potential candidate mechanism to generate clearing CSF flow is the recently discovered displacement of the LV borders induced by changes in total cerebral blood-volume (*14*). We hypothesized that – induced by cortical blood-volume changes - LV border movements are associated with LV-volume changes, which in turn drive CSF flow and clearance.

### Intracranial factors inducing LV-volume oscillations

#### Cortical vasodilation

To study whether cortical blood-volume changes induce LV border movements and associated volume changes, we first defined a method to track LV-border movements as a correlate of LV-volume changes by using the partial volume effects on standard fMRI-signals at the border between the LVs and the surrounding parenchyma (parenchyma-fluid-interface, PFI). Given that inhaling CO_2_ induces global vasodilation and thereby brain parenchyma volume increases (*21*), we performed a hypercapnic challenge in awake healthy adults breathing medical air alone or enriched with 5% CO_2_ (**Fig. 1A**). High-resolution T2-weighted (T2w) MRI across conditions revealed increased cortical gray matter (cortical-GM) volume (**Fig. 1B;** T = -7.11, P < 0.001) and decreased LV-volume (T = 4.23, P < 0.01) during hypercapnia. For each subject, subtracting T2w-MRI acquired during normocapnia from those acquired during hypercapnia resulted in a rim of voxels with negative signal intensities at LV borders, indicating that parenchyma (low T2w signal) displaced CSF (high T2w signal) (**Fig. S1A, Movie S1**). Susceptibility-weighted MRI under normo- and hypercapnia excluded a confounding effect from ependymal blood vessels (**Fig. S1B**).

**Fig. 1.**
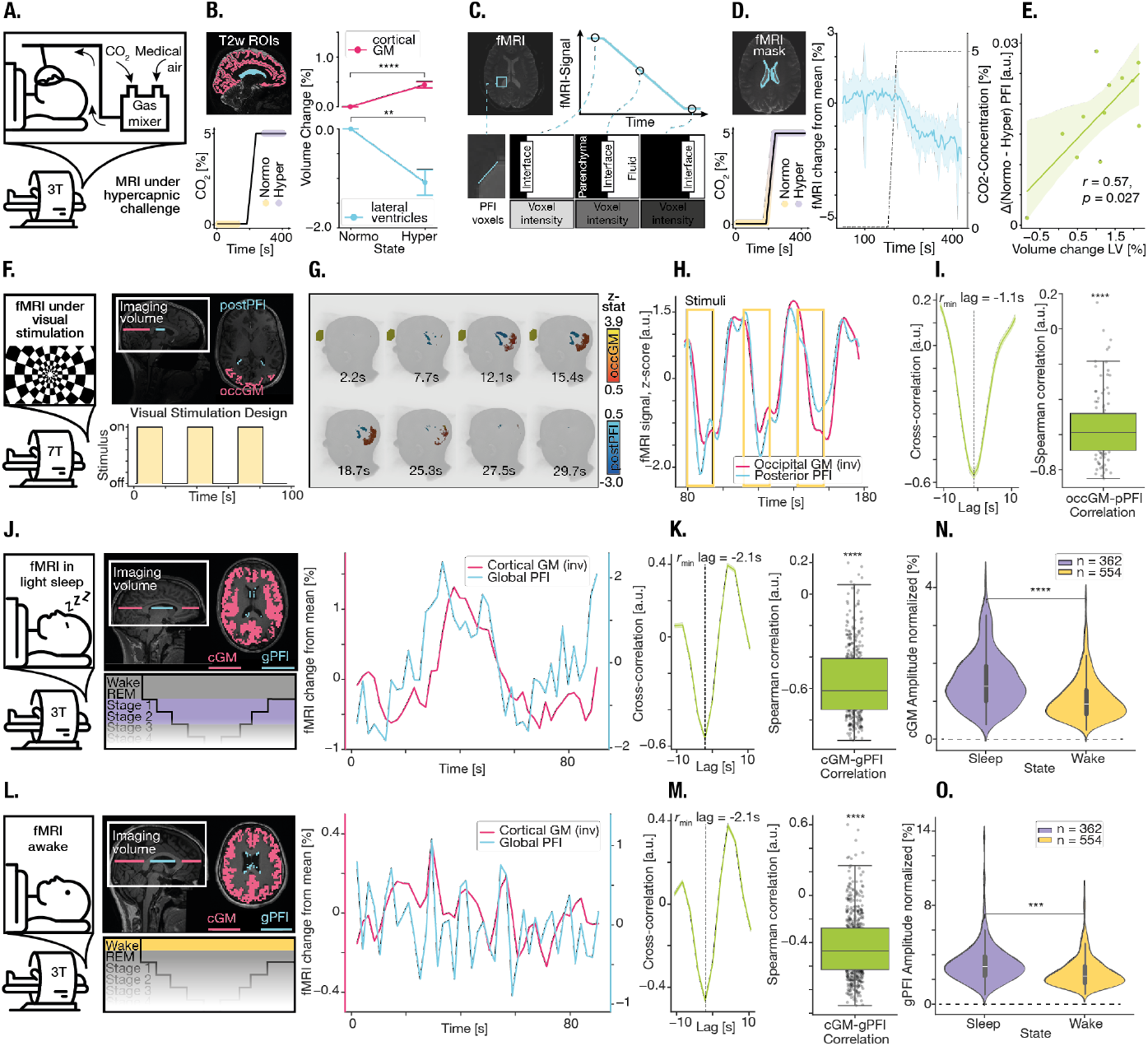
LV-volume changes during hypercapnia, visual stimulation, sleep and wakefulness. (**A**) Experimental setup. Healthy young subjects underwent an fMRI while breathing medical air alone or enriched with up to 5% CO_2_. (**B**) (*top left*): FreeSurfer-based segmentation of the LVs (*blue*) and cortex (*red*), superimposed on a sagittal T2 image from a representative subject. (*bottom left*): Timecourse of the hypercapnic challenge and time-point of T2 recordings (*shaded boxes*). 180s intervals of normo- or hypercapnia and 30s ramp time. T2 scans were acquired during stable normo- and hypercapnia (*shaded boxes*). (*right):* Normalized volume changes of the cortical GM (*red, top*) and LVs (*blue, bottom*) between normo- and hypercapnia. N= 12 subjects. Error bars depict SE. Paired t-test, t = -7.1, P < 0.0001, t = 4.2, P < 0.01. (**C**) Partial volume effects at the PFI. Due to the limited voxel size in the fMRI, the PFI is not fully resolved. The intensity of a respective PFI voxel (left) is determined by the ratio of CSF (high intensity) to parenchyma (low intensity). This ratio changes during the normo-hypercapnia transition, with CSF being replaced by tissue, resulting in a signal drop in these voxels (right). (**D**) (*top left*): segmented PFI mask at the LV edges (*blue*), superimposed on an axial fMRI image. (*bottom left*): timecourse of the hypercapnic challenge. fMRI recordings were performed continuously over the entire session, including stable baseline, transition and stable hypercapnia periods (*shaded box*). (*right*): Normalized global PFI timecourse (mean and SEM). N=12 subjects. The dotted line indicates the inhaled CO_2_ concentration. (**E**) Scatter plot of the LV-volume change during the first hypercapnic challenge and the PFI signal change in the second hypercapnic challenge. Each dot corresponds to one subject (N=12). Spearman rank correlation with one-sided t-test. (**F**) Experimental setup. Healthy young subjects underwent an fMRI and simultaneous visual stimulation with a flickering checkerboard (flicker frequency 12 Hz) in a block design (bottom). *Top right*: Placement of the masks for occGM (*red*) and posterior PFI (*blue*), superimposed on a T1 MPRage image. **(G)** 3D-rendering of the occGM and postPFI at different time points. The yellow box indicates the time interval of visual stimulation. (**H**) Mean timecourse of the occGM signal, inverted for visualization purposes (*red*), and the postPFI (*blue*) signal from a representative subject. The time of flicker stimulation is indicated in yellow boxes. (**I**) (*left*): Cross-correlation between cGM and gPFI for all subjects. mean and SEM (*shaded area*). The lag is defined as the point of highest anticorrelation between both signals. (*right*): Spearman correlation between cGM and gPFI. Each dot corresponds to one subject. One-sample t-test, t = -29.3, P < 0.0001. (**J**) Experimental setup. Healthy young subjects underwent an fMRI while sleeping. (*bottom*) 90s timeframes of NREM stage 1 and 2 were used. (*top right*): Placement of the masks for cGM (red) and global PFI (*blue*), superimposed on a T1 MPRage image. (*right*): Mean timecourse of the cGM signal, inverted for visualization purposes (*red*), and the globalPFI (*blue*) signal from a representative subject in sleep. (**K**) See Panel I, One-sample t-test, t = -56.6, P < 0.0001. (**L**) See Panel J, representative subject in wakefulness. (**M**) See Panel K, One-sample t-test, t = -38.6, P < 0.0001. (**N&O**) Higher cGM (N)- and gPFI (O)-amplitude during sleep than during wakefulness, 90s time-intervals N = 362 and N = 554, respectively, Mixed Linear Model, z = -3.9, P < 0,0001, z = -6.2, P < 0.0001.

To detect fMRI-signal intensity changes at the border of the LVs, we next defined the PFI mask as the rim of voxels generated by subtracting an LV mask eroded by one voxel in all three dimensions from the original LV-mask (**Fig. 1C, D & S1C**). Continuous fMRI-signals extracted from this mask revealed stable PFI signals under normocapnia, continuously decreasing signals during transitions from normo-to hypercapnia, and stable intensities under hypercapnia (**Fig. 1D**). While decreasing transition-related fMRI-signals indicated LV border movement because of changing PFI partial volume effects, lower median PFI signals during hyper-than normocapnia (T = 8.15, P < 0.001; **Fig. S1D**) demonstrated replacement of CSF (high T2w signal) by parenchyma (low T2w signal), indicating a decreased LV-volume under hypercapnia. Indeed, within subjects, the PFI fMRI signal decrease from normo-to hypercapnia was correlated with the magnitude of the anatomically determined LV-volume decrease (**Fig. 1E;** Spearman r = 0.57, P = 0.027), demonstrating that the PFI fMRI signal quantitatively tracks LV-volume changes induced by cortical vasodilation.

#### Visual stimulation

Next, we investigated whether experimentally induced cortical neuronal activity leads to LV-volume changes. Analyzing 7T fMRI during block-wise visual flicker stimulation (*22*), we confirmed that occipital-GM fMRI-signals, which reflect hemodynamics including blood-volume changes (*23*), increased after visual stimulation onset, remained stable during stimulation and decreased during no-stimulation periods (**Fig. 1F-I; Movie S2**). The fMRI-signals at the adjacent posterior part of the PFI decreased with stimulation onset and increased after stimulation end. Lag analysis revealed a delay of about 1s of occipital-GM signals relative to posterior-PFI signals (minimal r = -0.57 at lag -1.1s) with strong anti-correlations between both signals (mean r = – 0.56 +/- 0.195, P < 0.001). The delay of the cortical-GM relative to the PFI signal is likely due to the blood oxygenation level dependent (BOLD) delay (i.e., 2-3s) of cortical fMRI responses on neuronal stimulation (*23*) in contrast to PFI fMRI-signals which are more sensitive to changes in volume than blood oxygenation (*14, 24*), therefore reacting without delay. In contrast, regionally inconsistent GM/PFI signal pairs were less anti-correlated than occipital-GM/posterior-PFI (**Fig. S2B**; occGM/antPFI vs occGM/postPFI meandiff = -0.43, P < 0.001, prefrontGM/postPFI vs occGM/postPFI meandiff = -0.49, P < 0.001), indicating that localized cortical activity exclusively drives movement of the adjacent parts of the LV border. Finally, correlations between occipital-GM and ventricular CSF (without at-border voxels) were lower than occipital-GM/posterior-PFI anti-correlations (**Fig. S2B;** meandiff = -0.58, P < 0.001), excluding strong within-ventricle CSF effects on GM-induced PFI changes at ventricle borders.

#### Sleep and wakefulness

To test whether cortical-GM activity also influences LV-volumes under no-task conditions, we next analyzed resting-state EEG-fMRI data of adults during sleep and wakefulness (*25, 26*). We derived cortical activity and LV-volume oscillations via cortical-GM and global-PFI fMRI-signals, respectively, from subjects’ annotated periods of wakefulness and sleep (**Fig. 1J, L**). During sleep, lag analysis revealed a delay of cortical-GM signals relative to global-PFI (minimal r = -0.50 at lag -2.1s), with moderate anti-correlations between both signals (**Fig. 1K;** mean r = –0.48 +/- 0.258, P < 0.001), demonstrating induced LV-volume oscillations during sleep. During wakefulness, cortical-GM signals followed PFI signals (minimal r = -0.46 at -2.1s) and were moderately anticorrelated (**Fig. 1M;** mean r = – 0.43 +/- 0.263, P < 0.001), indicating induced LV-volume oscillations also for wakefulness. In addition, we observed larger amplitudes of cortical-GM (**Fig. 1N;** z = -6.21, P < 0.001) and PFI signals (**Fig. 1O**; z = -3.87, P < 0.001), as well as stronger cortical-GM/global-PFI anti-correlations during sleep than wakefulness (**Fig. S3A;** z = 5.10, P < 0.001), likely reflecting more coherent neuronal activity during sleep (*27*).

### Macroscopic CSF flow driven by LV-volume oscillations

Having demonstrated the impact of cortical blood-volume on LV-volume changes, we investigated next whether LV-volume changes induce CSF flow in the ventricular-cisternal pathway by mediating the impact of cortical blood and CSF-volume changes. We extracted fMRI-signals from masks of cortical-GM, PFI, and CSF at the bottom slice (bsCSF) of the imaging volume (**Fig. 2A**) for visual stimulation, sleep and wakefulness. BsCSF fMRI-signals reflect fluid entering the imaging volume (i.e., inflow effect) (*12, 28*). Correlations between the rate-of-change (i.e., the first derivative d/dt) of cortical-GM fMRI-signals and bsCSF fMRI signals reflect cortical blood-volume effects on CSF flow (*12, 14*). Relating the bsCSF-fMRI to the PFI and GM signals under visual stimulation (**Fig. 2B**), we observed that at the end of each stimulation block, sharp bsCSF peaks, indicative for CSF inflow, were associated with peaks of both the inverted (i.e., negative) d/dt occipital-GM, concurrent with a rapid drop of the occipital BOLD signal, as well as the d/dt posterior-PFI signal, indicative for high rates of increasing LV-volume. This observation suggests that expanding LVs are associated with CSF inward movement. Lag analysis revealed a delay of bsCSF signals relative to d/dt posterior-PFI signals (maximum r = 0.30 at lag -2.2s), with moderate correlations (**Fig. 2C;** mean r = 0.32 +/- 0.18, P < 0.001). Remarkably, the association between d/dt occipital-GM activity and bsCSF signals was mediated by d/dt posterior-PFI signals (**Fig. 2D, Table S1**; ab = -8.48, P < 0.0001), indicating that posterior LV-volume changes mechanistically translate the impact of occipital-GM blood-volume changes on CSF flow.

**Fig. 2.**
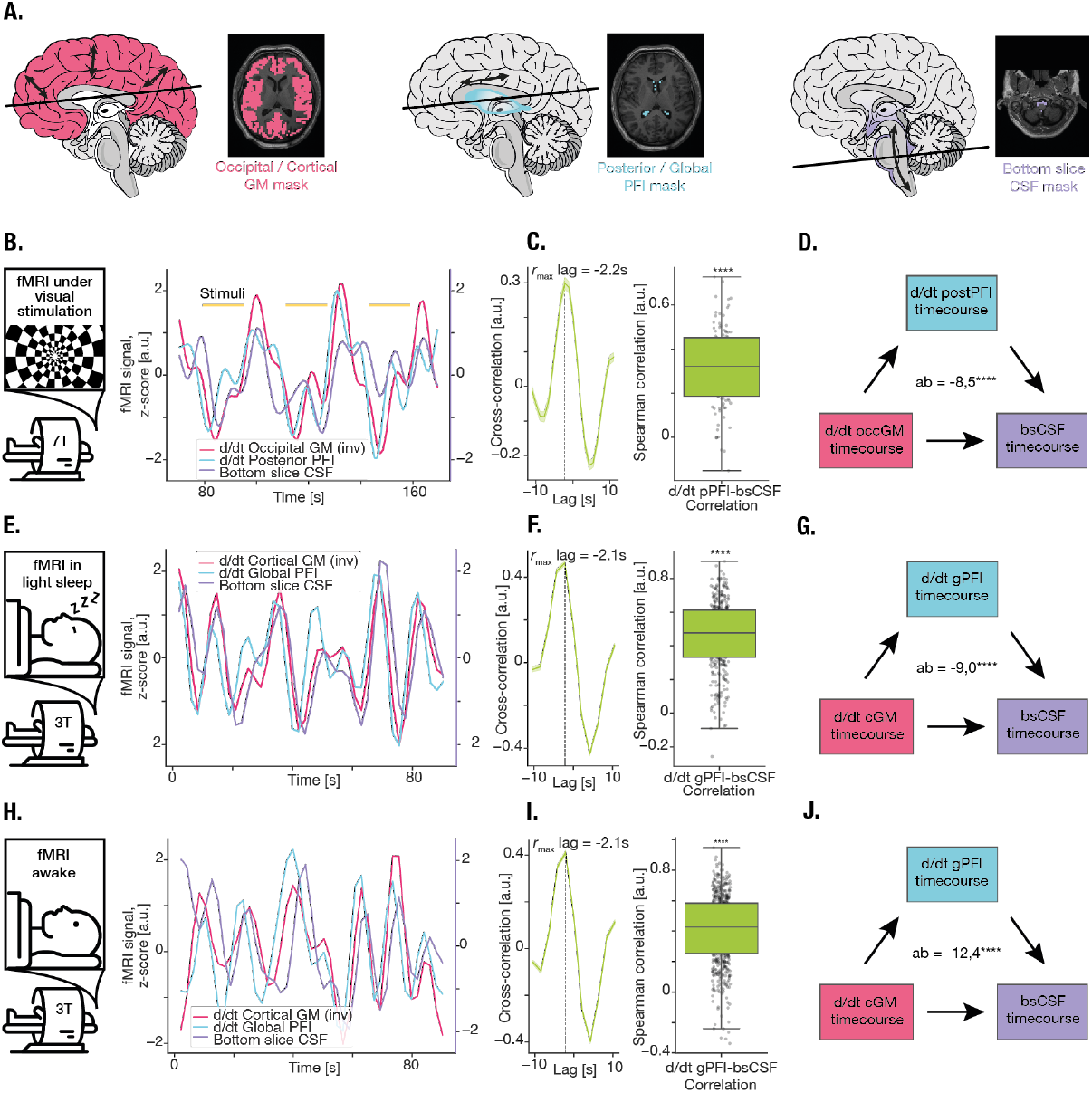
Undulating CSF flow induced and mediated by LV-volume oscillations. (**A**) Schematic of cortical GM, PFI, and ventricular-cisternal CSF imaging with exemplary masks superimposed on a T1 MPRage image from a representative subject. (**B**) During visual stimulation: Z-scored mean timecourses of occGM, postPFI and bsCSF from a representative subject. Stimuli are indicated in yellow. (**C**) Cross-correlation between occGM and postPFI for all subjects. Mean and SEM (*shaded area*). The lag is defined as the point of highest correlation between both signals. (*right)*: Spearman correlation between occGM and postPFI. Each dot corresponds to one subject. One-sample t-test, t = 18.7, P < 0.0001. (**D**) Multi-level mediation analysis between d/dt occGM, d/dt postPFI and bsCSF, shifted based on the results of the cross-correlation analyses shows a highly significant indirect path. ab = -8.48, P < 0.0001. (**E**) During sleep: Z-scored mean timecourses of cGM, gPFI and bsCSF from a representative subject in sleep. (**F**) Cross-correlation between cGM and gPFI for all subjects. mean and SEM (*shaded area*). The lag is defined as the point of highest correlation between both signals. (*right)*: Spearman correlation between cGM and gPFI. Each dot corresponds to one subject. One-sample t-test, t = 59.8, P < 0.0001. (**G**) Multi-level mediation analysis between d/dt cGM, d/dt gPFI and bsCSF, shifted based on the result of the cross-correlation analysis shows a highly significant indirect path, ab = -9.28, P < 0.0001. (**H**) During wakefulness: Z-scored mean timecourses of cGM, gPFI and bsCSF from a representative subject in wakefulness. (**I**) See Panel F, One-sample t-test, t = 42.9, P < 0.0001. (**J**) See Panel G, ab = -12.59, P < 0.0001.

During sleep and wakefulness (**Fig. 2E, J**), we observed similar inflow peaks in bsCSF associated with peaks in the -d/dt cortical-GM and the d/dt global-PFI signal. Both, lag and mediation analysis revealed a delay of the bsCSF signals relative to d/dt global-PFI signal (**Fig. 2F, I;** sleep: maximum r = 0.467 at lag -2.1s; for wakefulness: maximum r = 0.412 at lag -2.1s) with moderate correlations (sleep: mean r = 0.43 +/- 0.22, P < 0.001; wakefulness: mean r = 0.41 +/- 0.23, P < 0.001), as well as significant mediations of the associations between d/dt cortical GM and bsCSF by d/dt PFI signals for sleep (**Fig. 2G, J;** ab = -9.28, P < 0.0001) and wakefulness (ab = -12.59, P < 0.0001). Correlations between d/dt global-PFI and bsCSF signals were stronger in sleep than wakefulness (**Fig. S3B;** z = -3.13, P < 0.01), indicating stronger coupling between LV-volume oscillations and CSF flow during sleep. Thus, across largely different conditions, LV-volume changes drive downstream CSF flow with a delay of about 2.1s and mechanistically mediate the impact of cortical blood-volume changes on CSF flow (*12, 14, 20*).

### Extracranial factors inducing LV-volume oscillations

We asked next whether extracranial heartbeat and respiration (*15, 16, 18*) are also associated with LV-volume oscillations and its mediating impact on CSF flow. Using high temporal resolution 7T-fMRI data (TR = 227ms) and simultaneously registered cardiac and respiratory activity during resting wakefulness (*29*), we locked bandpass-filtered and Hilbert-transformed binned phase representations of cardiac and respiratory cycles to cortical-GM, global-PFI, and bsCSF traces, respectively (**Fig. 3A, G**). The d/dt PFI signal showed a significant oscillation along the cardiac cycle with a peak in the first and a trough in the second cycle half (**Fig. 3B**; trough = -0.50 %/bin, peak = 0.38 %/bin, median amplitude = 1.09%/bin, P < 0.0001). The d/dt cortical-GM signal also oscillates across the cardiac cycle but anti-correlated to the PFI signal changes, indicating an opposite link between changes in cortical blood- and LV-volumes during heartbeats (**Fig. 3C, D**; median amplitude = 0.30 %/bin, P < 0.0001; lag analysis: delay of d/dt PFI relative to d/dt cortical GM at 320° lag, with strong anti-correlations mean r = -0.73 +/- 0.27, P < 0.0001). In contrast, the bsCSF signal oscillates in parallel to the cardiac-induced PFI oscillations, indicating a direct mediating effect of LV-volume oscillations on CSF flow during heartbeats (**Fig. 3E, F;** trough = - 6.86 %, peak = 8.74 %, median amplitude = 19.77 %, P < 0.05; lag analysis: no delay of bsCSF relative to d/dt PFI, maximum r = 0.64 at 0° lag; correlations mean r = 0.64 +/- 0.25, P < 0.0001). Together, these results imply that during the diastolic filling of the heart, cortical blood-volume is low, leading to an expansion of the LVs and an inflow of CSF, while during the systole, blood-volume increases leading to a compression of the LVs and, putatively, to CSF outflow.

**Fig. 3.**
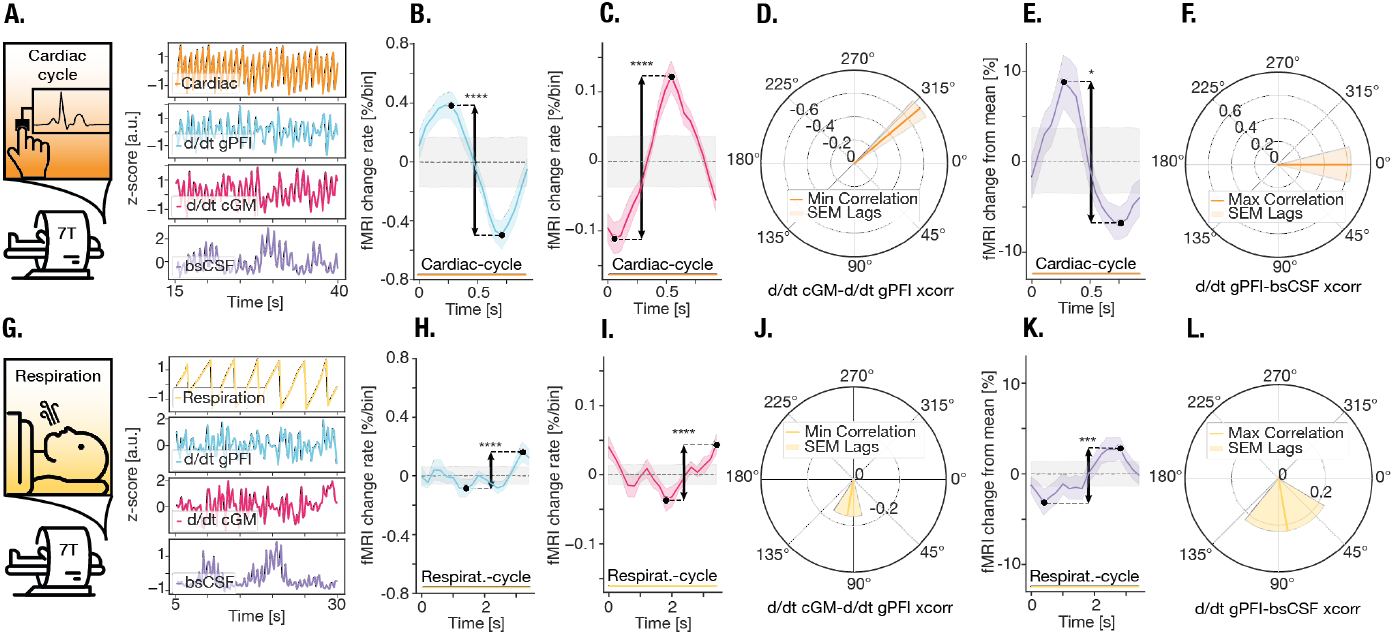
LV-volume oscillations induced by heartbeat and respiration. (**A**) (*left*): Experimental setup. Healthy young subjects underwent a 7T-fMRI while cardiac activity was recorded. (*right*): Z-scored cardiac data and mean d/dt cGM, d/dt gPFI and bsCSF timecourses from a representative subject. (**B**) Mean response of d/dt gPFI in each bin for all subjects to cardiac cycle, mean and SEM (*shaded coloured area*), permutated null distribution (*shaded gray area*). Peak-trough amplitudes were compared against the null distribution, Mann-Whitney U test, P < 0.0001. (**C**) same as B for d/dt cGM, P < 0.0001. (**D**) Cross-correlation between d/dt gPFI and d/dt cGM response for cardiac cycle for all subjects. mean and SEM (*shaded area*); delay of d/dt PFI relative to d/dt cGM, minimum r = - 0.73 at 320° lag. (**E**) same as B for d/dt bsCSF P < 0.05. (**F**) Cross-correlation between d/dt gPFI and bsCSF response for cardiac cycle for all subjects. mean and SEM (*shaded area*); no delay of d/dt PFI relative to bsCSF, maximum r = 0.64 at 0° lag. (**G - L**) For respiratory activity see Panel A-F, respectively.

Respiratory activity has, in terms of amplitude and oscillatory form, a weaker but still significant effect on LV and cortical blood-volumes and CSF flow (**Fig. 3G-L**). Respiratory-locked d/dt global-PFI, d/dt cortical-GM, and bsCSF signals fluctuate across the respiratory cycle (d/dt PFI/d/dt cortical GM/bsCSF: median amplitude = 0.35 %/bin, 0.11 %/bin, 11.08 %, P < 0.0001, P < 0.0001, P < 0.0001), and the d/dt PFI signal links with both, d/dt cortical-GM and bsCSF (delay relative to d/dt cortical GM at 100° lag with anti-correlation mean r = -0.16 +/- 0.15, P < 0.01, delay relative to bsCSF at 80° lag with correlation mean r = 0.26 +/- 0.26, P < 0.01). The weaker and less regular impact of respiratory activity on LV and cortical blood-volumes in comparison to cardiac activity might be due to distinct mediating pathways: intracranial pressure and venous volume changes for respiration, arterial volume changes for cardiac activity (*11, 15*).

### PET-tracer clearance of CSF driven by LV-volume oscillations

Having demonstrated their impact on CSF flow, we next studied whether LV-volume oscillations also drive substance clearance from the CSF. We performed simultaneous PET-fMRI in awake adults at-rest (**Fig. 4A)**. To estimate substance clearance, we measured the rate of ^18^F-DOPA-tracer decrease in the LVs by fitting linear regressions of LV tracer standardized uptake-value-ratios (SUVr) during dynamic PET (20-65 minutes), controlled for tracer decay (**Fig. 4B**); their slopes indicate the removal, i.e., ‘clearance’ of exogenously applied substances from the LVs’ CSF (*30*– *33*). ^18^F-DOPA-SUVRs monotonously decreased in all subjects (**Fig. 4C;** PET-tracer slopes: mean = -0.018 +/- 0.006), indicating consistent LV tracer clearance. The ventricular PET-tracer slopes did not correlate with mean nearby striatal ^18^F-DOPA-SUVr, ruling out a major impact derived from tracer spill-over from surrounding tissue (**Fig. S4A;** r = –0.05, P = 0.72).

**Fig. 4.**
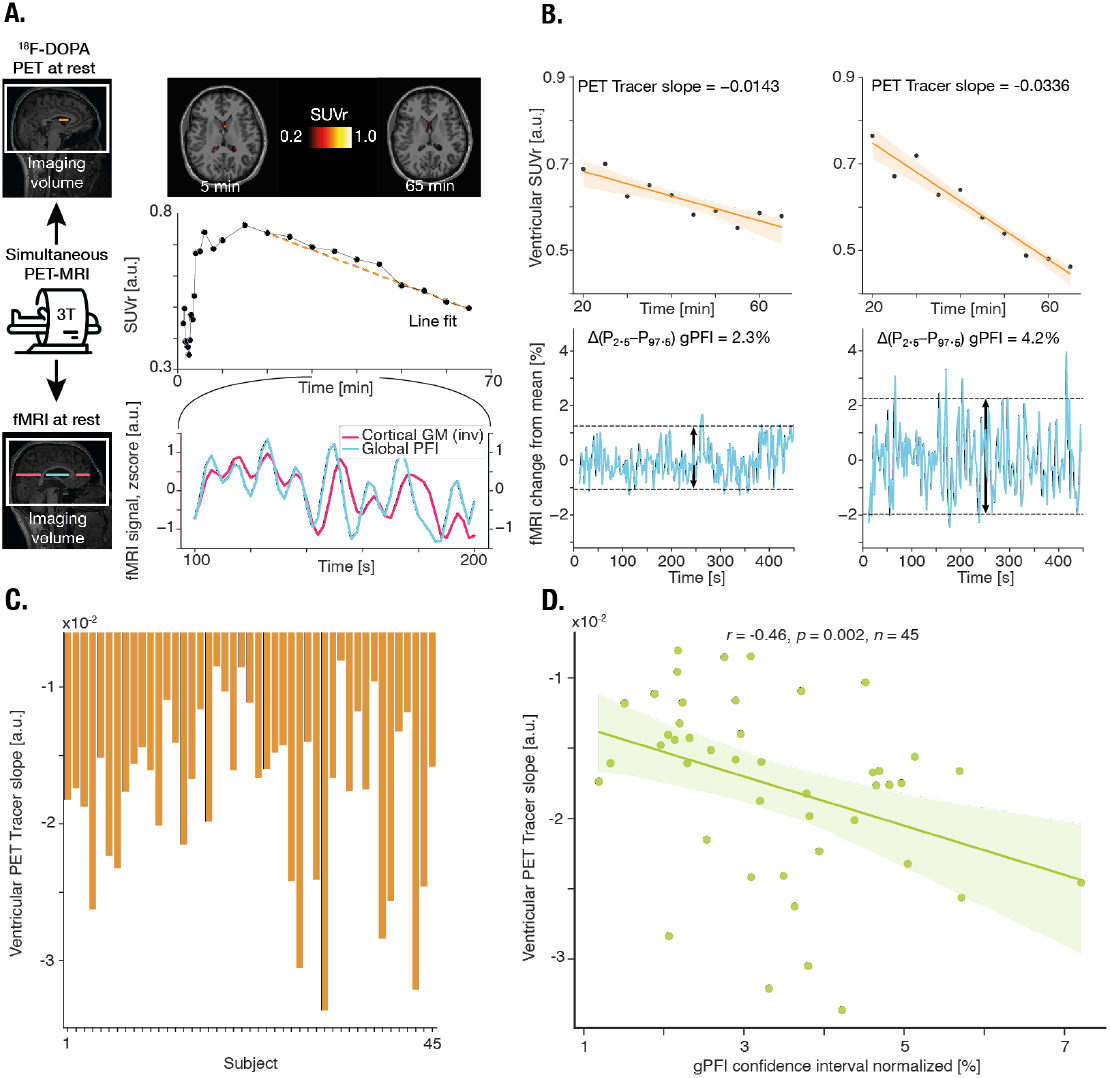
LV-volume oscillations and PET-tracer clearance. (**A**) Experimental design. Healthy young subjects underwent a simultaneous fMRI-^18^F-DOPA-PET scan. (*top right*): Mean lateral ventricular SUVr from a representative subject with transposed line fit between 20-65min to obtain regression slope (%/5min). (*bottom right*): z-scored mean d/dt gPFI and bsCSF timecourses from a representative subject. (**B**) Ventricular SUVr line fits of the eroded lateral ventricle mask and corresponding mean gPFI timecourses with calculated amplitudes (95%-confidence interval size) of two representative subjects. (**C**) Ventricular PET tracer slopes of all subjects. (**D**) Spearman rank correlation between ventricular PET tracer slope and gPFI amplitude. Each dot corresponds to one subject (N = 45), Spearman r = -0.46, P < 0.01.

In the simultaneously acquired fMRI data (28-43 min after tracer injection), we replicated the results of **Fig. 2** for simultaneously recorded cortical-GM, global-PFI, and bsCSF fMRI-signals, confirming that cortical blood-volume changes induce LV oscillations, which in turn mediate blood-volume impact on cisternal CSF flow (**Fig. S4B, C** lag analysis for cortical-GM and global-PFI fMRI-signals: minimum r = -0.52 at lag -2.0s; mean r = -0.50 +/- 0.21, P < 0.001; **Fig. S4D-F** lag and mediation analysis for d/dt cortical-GM, d/dt global-PFI, and bsCSF fMRI-signals: minimal r = 0.40 at -4.0s, mean r = 0.30 +/- 0.20, P < 0.001; ab = -3.64, P < 0.001). Then, we estimated the magnitude of LV-volume oscillations, as the 95%-confidence interval-based amplitude of the PFI signals (**Fig. 4B**), and linked them to the PET-tracer slopes of LVs. Remarkably, larger amplitudes of LV-volume oscillations were associated with a steeper decline of PET-tracer concentrations in the LVs (**Fig. 4D;** r = -0.46, P < 0.005), demonstrating that LV-volume oscillations indeed induce greater tracer clearance from the LVs’ CSF. In contrast, ventricular ^18^F-DOPA-slopes did not correlate with either the amplitudes of cortical blood-volume changes (i.e., amplitude of the cortical fMRI-signal; **Fig. S4G;** r = 0.24, P = 0.12) or the impact of cortical blood-volume changes on CSF flow (i.e., -d/dt cortical GM/bsCSF correlation coefficients; **Fig. S4H;** r = -0.19, P = 0.21), indicating that volume oscillations of the LVs but not of the cortex determine tracer clearance out of the ventricles.

### LV-volume oscillations and cortical blood-volume effects on CSF flow

In overall summary, these results demonstrate that LV-volume oscillations lead to undulating macroscopic CSF displacements in the ventricular-cisternal CSF system, which drive CSF substance clearance. LV-volume oscillations, in turn, are induced by distinct physiological drivers such as neuronal activity, heartbeat, and - to a lesser extent - respiration and mediated by cortical blood-volume changes.

While previous studies have reported an association between changes in global gray matter blood-volume and undulating CSF flow (*12, 14, 19*), the mechanistic drivers of this flow remained unclear. The mediation of the effect of cortical blood-volume oscillations on undulating CSF flow by LV-volume oscillations across different physiological conditions, scanners and MRI-modalities closes this gap by supporting a surprisingly simple model: blood-volume-induced balloon-like oscillatory expansion of the brain parenchyma displaces CSF in the ventricles, inducing macroscopic CSF flow. Furthermore, expanding previous findings of neuronal activity-dependent LV-volume change during burst-suppression anesthesia (*14*), our result generalizes this model as a common pathway of CSF flow generation for different physiological drivers.

On a microscopic scale, cortical arteriole diameter changes and associated periarteriole CSF bulk flow are induced by the same physiological drivers as macroscopic LV-volume changes and associated ventricular-cisternal CSF flow (*9*). Thus, induced arteriole blood-volume increases appear to integrate across scales to cortical blood and tissue volume increases with potentially complementary influences on CSF movement at micro-(i.e., periarteriole CSF) and macro-scale (i.e., ventricular-subarachnoid CSF). Our results suggest two potential pathways linking the macroscopic effects of physiological drivers on CSF with microscopic periarteriole CSF movements. Cortical blood-volume oscillations induce an expansion towards the surrounding external subarachnoid spaces (*14*), where CSF enters the cortical periarterial spaces before mixing with interstitial fluid (*2*). The expansion-associated volume changes of the external subarachnoid spaces may then induce further impact on periarteriole CSF movement beyond that of the physiological drivers on arteriole diameter (*34, 35*).

Additionally, subarachnoid cisternal CSF undulations connect with periarterial spaces of major cerebral artery trunks spreading from the basal cisternae to the whole-brain surface within external subarachnoid space. A recent tracer MRI study in humans demonstrated that gadobutrol-derived tracers spread from the basal cisternae directly into the periarterial spaces of these large cerebral arteries (*36*). Thus, LV-volume oscillations might drive this tracer spread along periarterial spaces eventually to the cortical periarteriole spaces.

### LV-volume oscillations and CSF substance clearance

The correlation between LV-volume oscillations and PET tracer decreases in LVs demonstrates a higher substance clearance from ventricular CSF in subjects with high LV oscillation amplitude, linking LV oscillations to brain clearance. So far, direct experimental evidence of macroscopic CSF flow being associated with clearance of substrates from the brain tissue or CSF is lacking due to limited options for measuring clearance of substances from the human brain. Besides the invasive intrathecal application of paramagnetic tracer in patient groups (*37, 38*), the PET-based approach used in the current study was recently suggested to track the clearance of a blood-brain-barrier permeant, i.v.-injected tracer from the CSF over time (*30*). After its initial deposition in the CSF via the choroid plexus when blood tracer activity is high, tracer activity linearly drops over time indicating gradual replacement of CSF by CSF with lower tracer concentrations (*31*–*33, 39*). This is likely due to the production of ‘fresh’ CSF and/or by mixing with tracer-free CSF in the cisternal outflow paths. Due to the low rate of choroid plexus CSF production (*2, 40*) in comparison to our measurement duration, our result of CSF-tracer clearance correlating with the magnitude of LV-volume oscillations suggests that mixing processes are significantly enhanced by large undulating flow events of CSF. Furthermore, since periarterial spaces of large arteries are a major cisternal outflow path (*36, 41*), our finding supports the model that LV induced CSF oscillations drive substance movement along subarachnoid periarterial spaces.

Together, our data demonstrate that LV oscillations are a major driver of CSF flow and clearance in the human brain. In consequence, enhancing LV-oscillations may have beneficial effects on brain clearance and should be explored as a disease modifying strategy in conditions with impaired brain clearance.

## Author contributions

Conceptualization: VN, MB, BZ, CS

Methodology: VN, MB, GH, CP, FB, IY, BZ, CS

Investigation: VN, MB, MT, JS, FB

Visualization: VN, MB, BZ

Funding acquisition: FB, IY, CP, BZ, CS

Project administration: VN, MB, MT, JS

Supervision: BZ, CS

Writing – original draft: VN, MB, BZ, CS

Writing – review & editing: VN, MB, MT, JS, MB, JK, JZ, JP, GH, CP, FB, IY, BZ, CS

## Data and materials availability

Data from visual stimulation study, sleep study and extra-cranial physiological factors study can be found at the OpenNeuro database (https://openneuro.org/datasets/ds004484/versions/1.0.0, https://openneuro.org/datasets/ds003768/versions/1.0.10, https://openneuro.org/datasets/ds004645/versions/1.0.0).

In-house data from the hypercapnia and simultaneous ^18^F-DOPA-PET-fMRI study will be made available by the corresponding authors upon reasonable request.

All data tables and code used to perform the presented analyses can be found at https://github.com/VikNeu/LV-CSF-flow-clear

## Supplementary Materials

Materials and Methods

Supplementary Text

Figs. S1 to S4

Table S1

Movie S1, S2

## Materials and Methods

### Subjects, experiments, and data acquisition

Experimental data from three independent in-house studies and three independent open-access studies (i.e., from the OpenNeuro data base) of healthy adults, respectively, were used for the current study.

In-house studies were conducted in line with the Declaration of Helsinki and approved by the ethics committee of the medical school of the Technical University of Munich (institutional review board approval numbers 472/16S for the hypercapnia study and 5611/12 and 338/20S for the two PET-MRI experiments of the PET-fMRI study, respectively). Participants were given detailed information about the methods, potential risks, and gave their written informed consent before the experiments.

#### Hypercapnia study (Fig. 1A-E)

Fifteen healthy participants (mean age 24.1, 8 female) underwent 3T fMRI during a controlled hypercapnic challenge at TUM University Hospital. We applied three cycles of alternating medical air alone or enriched with 5% CO_2_ for ∼180s each (30 s ramp time) using a Gasmixer (AltiTrainer, SMTec, Switzerland) via a sealed face mask. Scans were acquired on 3T Philips Ingenia MR-Scanner using a 32-channel head coil. Structural MRI data were acquired using T1-weighted (T1w) magnetization prepared rapid gradient echo (MPRAGE; TE = 4ms, TR = 9ms, flip angle = 8 degrees, FOV = 240mm × 252mm × 200.25mm, voxel size = 0.75mm × 0.75mm × 0.75mm) and T2-weighted (T2w) Turbo Spin Echo (TSE; voxel size = 0.7 mm × 0.7mm × 0.7mm). Functional MRI data were acquired using single-shot gradient echo (GE) echo planar imaging (EPI; TE = 80ms, TR = 4100ms, number of slices = 20, FOV = 205mm × 205mm × 30mm, voxel size = 0.8mm × 0.8mm × 1.5mm). To visualize venous vasculature, we acquired gradient echo data for susceptibility weighted imaging (SWI; TE = 9.8ms, TR = 27ms, flip angle = 20 degrees, FOV = 230mm × 188.47mm × 141mm, voxel size = 0.8mm × 0.8mm × 1.5mm).

Three subjects were excluded due to a failure in the scanning protocol (N = 1) or excessive head motion (N = 2).

#### Visual stimulation study (Fig. 1F-I and 2B-D)

We used an open-access dataset (https://openneuro.org/datasets/ds004484/versions/1.0.0) from a previously published 7T fMRI study of 20 healthy participants (*20, 22*), who underwent visual stimulation in a block design during fMRI by viewing a radial checkerboard flickering at a rate of 12 Hz interspersed with equally long episodes of a gray screen (flicker blocks ranging from 0.176 to 16 s). All subjects gave written informed consent to procedures approved by Massachusetts General Hospital’s Institutional Review Board (IRB #2014P001038).

Scans were acquired on a 7T Siemens MAGNETOM scanner with a custom-built 32-channel head coil. For our analysis, we used the structural T1w MPRAGE (isotropic spatial resolution of 0.75mm^3^) and the functional MRI data acquired by simultaneous multi-slice (SMS) single-shot gradient echo EPI (isotropic spatial resolution of 1.1mm^3^, TE = 26 ms, TR = 1.11 s, 38 slices, R = 4 acceleration, Multiband factor = 2, matrix = 174 × 174 full-Fourier, blipped CAIPI shift = FOV/2, flip angle = 70°)(*20*).

We excluded one subject because spatial normalization by Talairach transformation failed.

#### Sleep study (Fig. 1J-O and 2E-J)

We used an open-access dataset (https://openneuro.org/datasets/ds003768/versions/1.0.10) from a previously published 3T study of 33 healthy participants, collected at Pennsylvania State University (*25, 26*). In the original study, subjects underwent anatomical MRI, followed by two 10-minute awake resting-state and multiple 15-minute sleep scans on a 3T Prisma Siemens Fit scanner with a Siemens 20-channel receive-array coil, while EEG data was simultaneously acquired using an fMRI-compatible 32-channel electrode cap (BrainAmp MR, Brain Products, Germany). For our analysis, we used the anatomical MPRAGE (TE = 2.28ms, TR = 2300ms, 1mm isotropic spatial resolution, FOV = 256 millimeters, flip angle = 8 degrees, matrix size = 256×256×192, acceleration factor = 2) and EPI data (TE = 25 ms, TR = 2100 ms, slices = 35, FOV = 240mm, voxel size = 3mm × 3mm × 4mm), as well as the pre-labelled EEG data (*25, 26*). Based on the classification of the subjects’ arousal state into wakefulness (‘Wake’) and non-rapid eye movement (NREM) stages 1, 2, and 3 (*26*), we analyzed time frames where the subject stayed in the same state (wake, NREM stage 1 or 2) for at least 150s. We omitted NREM 3, because subjects typically did not reach stable NREM 3. As suggested by the literature (*12*), the first and last 30s of the 150s intervals were cut to avoid data from transition states. That left us with 554 90s-timeframes in wakefulness and 362 90s-timeframes in NREM 1 and NREM 2 together across all subjects to analyze.

#### Extra-cranial physiological factors study (Fig. 3)

We used an open-access dataset (https://openneuro.org/datasets/ds004645/versions/1.0.0) from a previously published 7T-fMRI study (*29, 42*), where 15 subjects underwent ultrafast resting-state fMRI on a 7T Siemens MAGNETOM scanner with a custom-built 32-channel head coil while their heart rate and respiration were monitored using a piezoelectric transducer and a respiratory belt at a sampling rate of 1000 Hz.

For our analysis, we used the structural multi-echo magnetization-prepared rapid gradient-echo (MEMPRAGE; isotropic spatial resolution of 0.75mm^3^, TE =1.76 ms and 3.7ms, echo-spacing=6.2ms, TR = 2.530ms, TI = 1100ms, 7° flip angle, bandwidth = 651 Hz, in-plane acceleration R=2, FOV = 320 × 320 × 244 mm, total scan time of 7:20 min) and the functional MRI single-shot GE EPI data (2mm isotropic spatial resolution, TE = 24ms, echo-spacing=0.59ms, TR = 227ms, 30° flip angle, bandwidth = 2604 Hz, in-plane acceleration R=2, SMS Multiband Factor = 3, CAIPI shift = FOV/3) (*6*).

We excluded one subject because spatial normalization by Talairach transformation failed.

#### Simultaneous ^18^F-DOPA-PET-fMRI study (Fig. 4)

Data for this study were collected from two independent PET-fMRI experiments at the same hybrid 3T PET-MRI scanner (mMR Biograph, Siemens Healthcare) at TUM University Hospital, but with different MR head coils. One experiment used data from 22 healthy subjects who underwent simultaneous fMRI and ^18^F-DOPA PET using a 32-channel phased-array head coil. ^18^F-DOPA was intravenously injected 30s before the simultaneous PET/MRI started, with a total acquisition time of 70min for each participant. Ordered subset expectation maximization (21 subsets, 3 iterations) was used to reconstruct PET data with a voxel size of 1.7×1.7×2mm^3^, corrected for attenuation, decay, and scatter based on anatomical MRI information. PET data were framed into 30 dynamic frames (1×30s, 10×15s, 3×20s, 2×60s, 2×120s, 12×300s). Structural T1w MRI (MPRAGE; TR = 2300 ms, TE=2.98ms, TI = 900 ms, phase encoding direction = anterior to posterior, flip angle = 9 degrees, FOV = 256 mm, matrix size = 240 × 256 × 160, voxel size = 1 mm × 1 mm × 1 mm) was performed 10 minutes after PET acquisition started, rs-fMRI (GE EPI; TE = 30ms, TR = 2000ms, flip angle = 90 degrees, slice thickness = 3mm, number of slices = 35,

FOV = 192mm, in-plane resolution = 2mm × 2mm) acquisition began 28 minutes after PET start and lasted for 15 minutes. Before rs-fMRI, participants were instructed to close their eyes, remain still, and stay awake.

The second experiment contained data from 24 subjects, which has been published previously (*43*). Subjects underwent simultaneous fMRI and ^18^F-DOPA PET at the same hybrid whole-body mMR Biograph PET/MRI scanner, but using a 12-channel phased-array head coil. PET and structural MRI protocols were identical to the first experiment. For fMRI, GE EPI used different sequence parameters (TE=30ms, TR=2000ms, 90-degree flip angle, 35 axial slices, slice thickness=3.0mm, FOV=192×192mm, in-plane resolution = 3.0mmx3.0mm).

One subject of the first experiment was excluded because of a failure in the PET scan protocol.

### Brain MRI data preprocessing and brain mask creation

#### MRI data preprocessing

##### Structural MRI

T1-weighted MRI data from all experiments were preprocessed using recon-all from FreeSurfer 7.4.1 (*44*). T2-weighted MRI data from Experiment 1 were preprocessed using recon-all-clinical from FreeSurfer 7.4.1 (*45*).

##### Functional MRI

GE EPI data from all experiments were preprocessed using FEAT from FSL version 6.0.7.6 (*46, 47*). In each case, we used the longest time frame in the fMRI data with no field-wise displacement of over 2mm. After removal of the first 5 volumes, we performed motion correction with MCFLIRT, slice time correction for data with a TR > 1s, and denoising of physiological confounders with independent component analysis using ICA-Aroma (*48*). For ICA-AROMA the fMRI image was brain extracted using BET. No spatial smoothing was applied.

#### Mask creation for MRI

##### Cortical gray matter masks

Gray matter (GM) masks were created using the FSL fast command. The output generated was then intersected with the Harvard Oxford Cortical atlas, which was registered to subject space, to focus on cortical GM structures. In a last step, all voxels intersecting with the cerebellum of the MNI atlas in subject space, were subtracted. Occipital and prefrontal (i.e., superior-frontal) cortical gray matter masks (occGM, prefrontGM) were created based on the structural MRI using FreeSurfer (aseg+aparc file region 1011/2011 and 1028/2028, respectively) and then transferred into the fMRI space using the FSL flirt command.

##### PFI masks

For the lateral ventricular border (i.e., the parenchyma-fluid-interface, PFI) masks, lateral ventricle masks were created first on the structural MRI using FreeSurfer and then transferred into the fMRI space using the FSL flirt command. The masks were eroded by one voxel and subtracted from the lateral ventricle mask (Fig. S1). For our low-resolution data (fMRI data of the sleep and simultaneous ^18^F-DOPA-PET-fMRI studies, i.e., voxel dimensions > 2 mm), we excluded voxels that did not contain a fraction of CSF based on two conditions. Only voxels with, (I) on-average lower intensity during inflow events (CSF intensity z-score >1) than during outflow events (CSF intensity z-score <0) and (II) low correlation with the mean gray matter timecourse (Pearson’s r <0.1) were included in the mask. To create the regional PFI (rPFI) masks for the visual stimulation study (in contrast to the global-PFI (gPFI) of the whole lateral ventricles), the gPFI mask was intersected with a manual segmentation of the anterior or posterior border of the ventricle (antPFI, postPFI) that was created in MNI space and then transferred into subject space using the FSL flirt command.

##### CSF masks

Bottom-slice CSF (bsCSF) masks were created semi-automatically according to a previously published protocol (*14, 49*). In short, we first thresholded voxels that had high intensity, variance, and high maximum derivative due to the higher intensity of CSF compared to tissue in EPI sequences (inflow-effect). This first approximation was then corrected manually using ITK Snap (*50*), if needed.

#### PET data preprocessing

PET data were analyzed along with the T1-weighted MRI data using SPM12 (*51*). PET data were corrected for motion by realigning all PET frames to the mean time frame. Since anatomical information was not sufficient in the first frames, the transformation matrix of the frame at minute 5 was applied to all preceding frames. The individual T1w data was co-registered to the mean-time frame and then spatially normalized into MNI space. The inverse transformation matrix of this normalization was then applied to all regions-of-interest to transform them to the individual subject’s PET space. The standardized uptake value (SUV) was calculated for all dynamic PET time frames by normalizing the reconstructed activity by injected dose and body weight.

##### Mask creation for PET

Brain masks (lateral ventricles and striatum) were created based on the structural MRI using FreeSurfer and then transferred into the PET space using the FSL flirt command.

Eroded lateral ventricle masks were created by shrinking the lateral ventricle mask by 1 voxel (i.e., 3 mm^3^) to avoid any contamination of the lateral ventricular PET tracer signal from surrounding structures.

### Data analysis, outcomes and statistical analysis

Data analysis as well as statistical analysis were performed in Matlab21b and Python 3.13.2 using the Scipy (*52*), statsmodels (*53*) and pingouin (*54*) toolboxes. All plots were created with matplotlib (*55*) and seaborn (*56*). Color palette is Monokai Pro (*57*).

#### FMRI-outcomes and statistical analysis

For each subject, preprocessed fMRI timecourses were extracted from the respective masks (i.e., cortical GM, PFI, and CSF) using CoSMoMVPA software (*58*) for Matlab21b. After calculating the mean of all masked voxels for each mask, respectively, the timecourses were linearly detrended and bandpass-filtered between 0.01 and 0.1 Hz.

#### Amplitudes

PFI amplitude (of Figure 1 analysis) was calculated as the difference between the maximum and minimum of the PFI-fMRI timecourse and then normalized by dividing by the mean of the corresponding timecourse.

PFI amplitude (of Figure 4 analysis) as 95%-Confidence Interval: For each normalized and detrended timecourse, the 2.5th and 97.5th timecourse percentiles were calculated to capture the distributional spread and account for potential influence of movement on the PFI signal.

Median signal change from mean (Figure 1) was obtained from timeframe 25 to 44 in Normocapnia and 65 to 84 in Hypercapnia to guarantee steady states.

#### Z-scoring and change from mean

To obtain z-transformed timecourses, fMRI timecourses were z-scored (using zscore from SciPy 1.15.3). To obtain fMRI signal change from mean timecourses (S(t)/S_0_), fMRI timecourses without filtering were divided by the mean value of the timecourse.

#### Cross-correlation and correlation analysis

The lag between two fMRI timecourses was calculated by cross-correlation (xcorr function Matlab21b with a maximum lag of 15 TRs). Correlations were calculated using the Spearman rank correlation.

The coupling values of the group were tested against zero using a one-sample t-test (stats.ttest_1samp from SciPy 1.15.3).

#### Mediation analysis

Mediation analysis was performed using the mediation.m function of the canlab-MediationToolbox-19917e7 (https://github.com/canlab/MediationToolbox) for a multi-level mediation (*59*) of the d/dt GM, d/dt pPFI and bsCSF timecourses of all subjects (see Figure 2). d/dt GM and bsCSF timecourses were shifted forward following the results of the cross-correlations. The output consisted of model paths and their statistics, of which the indirect path was reported in the main text as evidence for a mediation effect. The complete output of each mediation can be found in Table S1.

#### Sample comparisons

For sample comparisons in both visual stimulation and sleep studies, where multiple events were drawn from the same subject, a Mixed Linear Model (statsmodels.mixedlm) was used to compare different states. For Fig. S2, to compare correlation coefficients of different GM/PFI signal pairs, we used Mixed Linear Model with pairwise tukey post hoc test, correcting for multiple comparisons (pairwise_tukeyhsd from statsmodels.multicomp).

#### Analysis of heartbeat and respiration data

For the extra-cranial physiological factor study, we processed the physiological cardiac and respiratory data that were recorded by using a piezoelectric transducer and a respiratory belt for each subject (*42*). These data were detrended and bandpass filtered (0.1-0.4 Hz for the respiratory traces and 0.6-1.3 Hz for the cardiac traces), following previous approaches (*20*). We then calculated the phase of the filtered physiological signals using the Hilbert transformation, separating 18 bins of 20 degrees, respectively. After that, we averaged the d/dt cGM, d/dt gPFI and bsCSF response, respectively, in each bin for every subject and then across subjects. To obtain signal change from the mean (S(t)/S_0_) similarly as described above, averaged responses were normalized to the timecourse mean. Regarding the first temporal derivatives of cGM and gPFI timecourses, signal change is given in %/bin, whilst bsCSF is given in %. The response plots B, C, E, H, I and K in Figure 3 indicate the mean and standard error of the d/dt cGM, d/dt gPFI and bsCSF response to the cardiac and respiratory cycle over subjects. To generate a null distribution for each physiological analysis (i.e., respiration and cardiac) for d/dt cGM, d/dt gPFI, and bsCSF, respectively, the bin indices used to extract the data points in the Hilbert transformation were randomly shuffled using Matlab’s randperm function. This procedure was repeated 10.000 times to establish a robust distribution of surrogate data. In the response plots of Figure 3, the lighter areas represent the standard error of the mean (SEM), and the darker lines show the mean signal of the first temporal derivative of cGM and gPFI as well as mean signal of bsCSF.

#### Statistical analysis

Trough to peak amplitude values (max-min) were statistically compared against the amplitudes of the confidence interval of the null distribution (see PFI amplitude Figure 4) using Mann–Whitney U test (mannwhitneyu from SciPy 1.15.3). To assess temporal lag between the first derivative of gPFI and bsCSF signals across the two physiological conditions, cross-correlations were computed for each subject using scipy.signal.correlate. The lag corresponding to the maximum (anti-)correlation between the two variables of interest was identified and subsequently transformed into angular values to match the bin structure of the analysis. Orange (cardiac cycle) and yellow lines (respiratory cycle) represent the mean angles of maximum correlation across subjects, while the shaded regions indicate the standard error of the mean (SEM) for these angles.

#### Ventricular PET tracer slope and its link with ventricular fMRI measures

For each subject, preprocessed PET images (SUV) were normalized to the intensity of the whole brain at 20 minutes (SUVr), to control for different amounts of PET-tracer being delivered to the brain (*30, 31*). Then a linear model (fitlm from Matlab21b) was fitted to the mean timecourse of voxels within the eroded lateral ventricle mask. The regression coefficient of this model was used as the rate of PET tracer clearance from the lateral ventricle (percentage decrease/5min). For the striatum, the mean value of SUVr within the striatal mask was calculated.

#### Correlation analysis

Correlation between ventricular PET tracer slope and normalized PFI confidence interval was calculated using the Spearman rank correlation.

## Tables

**Table S1.** Results of Multilevel mediation analysis assessing d/dt PFI as a mediator for the association between d/dt GM and bsCSF activity.

| Study: state | a | b | c' | c | ab |
| --- | --- | --- | --- | --- | --- |
| Visual stimulation | -19.61**** | 10.91**** | -2.70* | -10.85**** | -8.48**** |
| Sleep: sleep state | -34.27**** | 11.65**** | -27.79**** | -39.20**** | -9.28**** |
| Sleep: wake state | -31.42**** | 18.22**** | -27.66**** | -35.22**** | -12.59**** |
| PET-fMRI | -8.42**** | 5.16**** | -11.14**** | -11.64**** | -3.64*** |

**Note:** Results are expressed as t-values with two-tailed p-values.
\*\*\*\* $P \leq 0.0001$ \*\*\* $0.0001 < P \leq 0.001$ \*\* $0.001 < P \leq 0.01$ \* $0.01 < P \leq 0.05$ ns $P > 0.05$ (not significant)

## Movies

***Movie S1***. 3D-stack of the subtracted T2 image of the subject shown in Fig. S1A.

***Movie S2***. Animated movie showing the dynamic intensities of occGM and postPFI during visual stimulation.

