## Supplement Figures for "Cerebrospinal fluid flow and clearance driven by lateral ventricle volume oscillations"

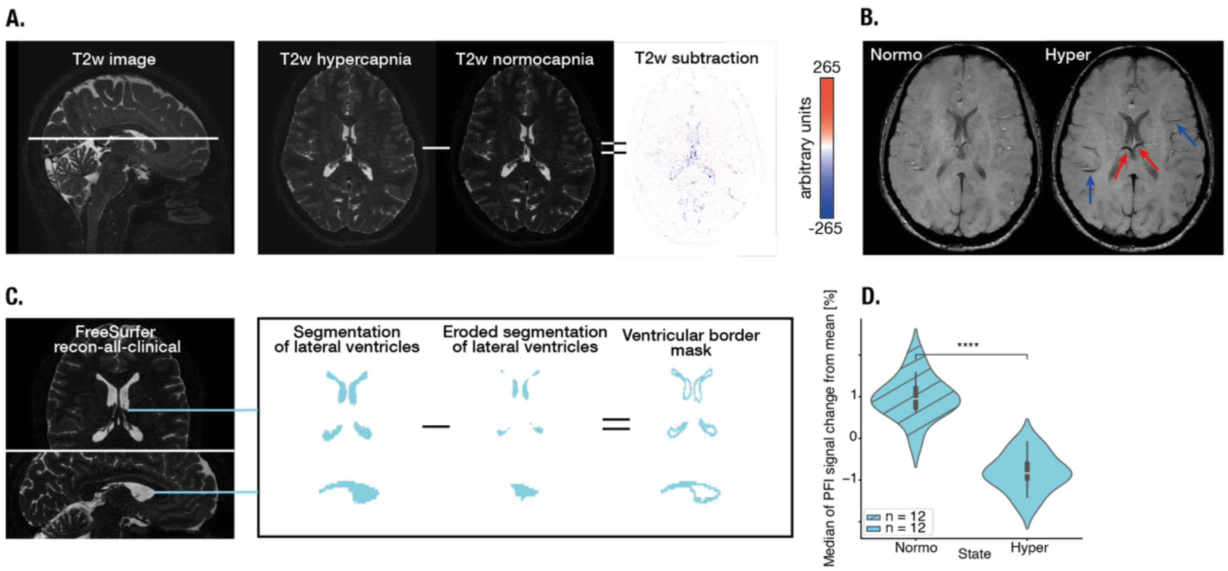


**Fig. S1. LVs are compressed by hypercapnia.**

(**A**) T2w images from a representative subject under normo- and hypercapnia and subtraction image (hypercapnia – normocapnia, *right*). Note the negative intensities in the subtraction image on the LV borders, indicating that during hypercapnia, tissue (low intensity) displaces CSF (high intensity). (**B**) Susceptibility weighted images (SWI), sensitive for blood vessels, appearing dark under normo- and hypercapnia. Blue arrows indicate outer and red arrows indicate inner cortical veins. (**C**) Generation of the ventricular border mask. (**D**) PFI signal intensities during normocapnia and hypercapnia. Paired t-test, $t = 8.15$, $P < 0.001$.

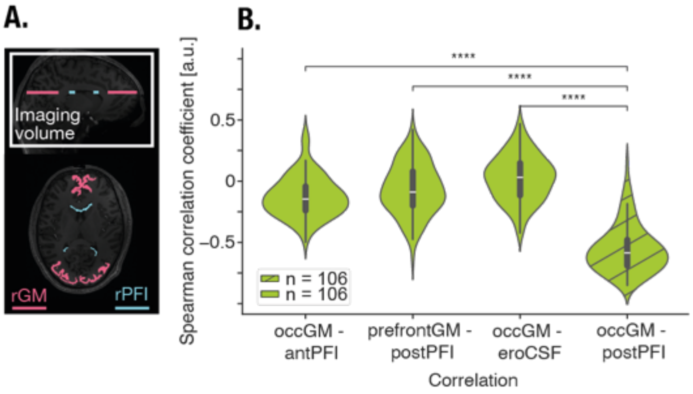


**Fig. S2. Correlations between regionally inconsistent GM-PFI pairs.**

(**A**) To control that only regionally consistent GM regions drive corresponding PFI regions, distinct GM and PFI regions have been defined: prefrontal and occipital GM (*red*), and anterior and posterior PFI (*blue*) masks, superimposed on a T1 image. (**B**) Comparisons of Spearman correlation coefficients of regionally inconsistent GM/PFI pairs occGM/antPFI, prefrontGM/postPFI, occGM/eroCSF, respectively, with consistent occGM/postPFI; eroCSF refers to an eroded LV mask without at-border voxels to control whether there is a correlation between occipital GM activity and a ventricular only CSF signal; Mixed Linear Model with pairwise tukey post hoc test, meandiff = -0.43, $P < 0.0001$ (*far left*), meandiff = -0.49, $P < 0.0001$ (*left),* meandiff = -0.58, $P < 0.0001$ (*right),* N = 106.

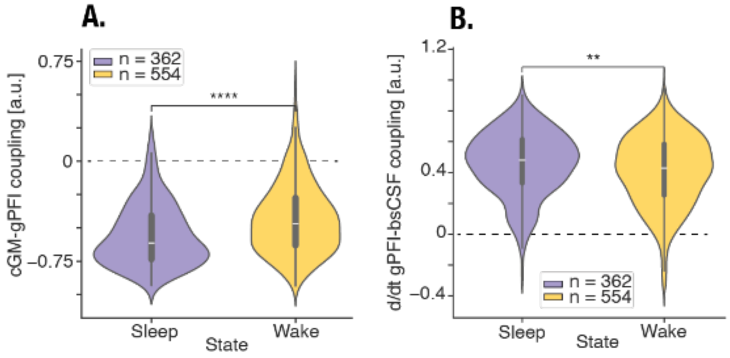


**Fig. S3. Stronger cGM-gPFI and gPFI-bsCSF coupling during sleep than during wakefulness.**

We identified 362 and 554 90s time-intervals for sleep and wakefulness, respectively, across 33 subjects during EEG-fMRI at resting-state. (**A**) The coupling between cGM and gPFI fMRI signal was stronger during sleep than wakefulness, tested using a Mixed Linear Model: $z = 5.1$, $P < 0.0001$. (**B**) The coupling between d/dt gPFI and bsCSF fMRI signal was stronger during sleep than wakefulness, tested by using a Mixed Linear Model: $z = -3.1$, $P < 0.01$

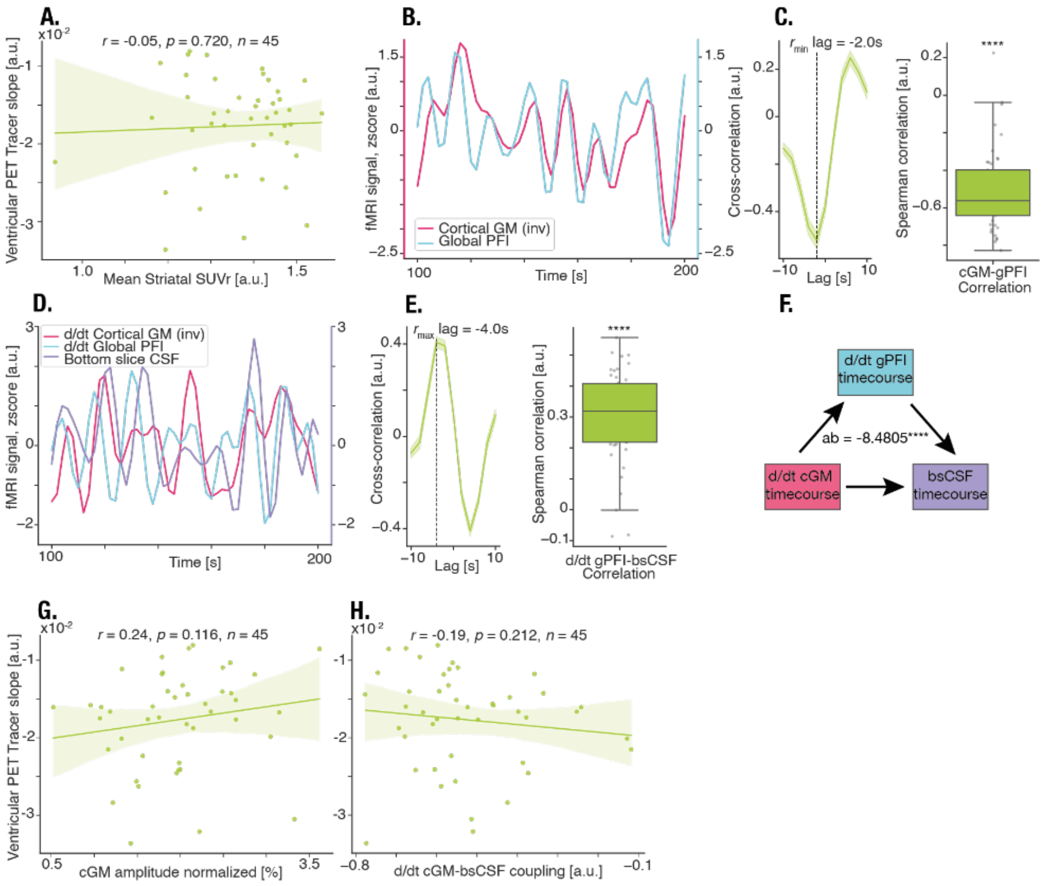


**Fig. S4. Control analyses for ventricular PET tracer clearance and its link with LV volume oscillations.**

(**A**) Spearman rank correlation between ventricular PET tracer slopes and striatal SUVrs. (**B**) z-scored mean time courses of cGM *(red*) and gPFI (*blue*) from a representative subject. (**C**) (*left):* Cross-correlation and spearman correlation between cGM and gPFI for all subjects, mean and SEM (*shaded area*). The lag is defined as the point of highest anticorrelation between both signals. (*right)*: Spearman correlation between cGM and gPFI. Each dot corresponds to one subject. One-sample t-test, t = -16.2, P < 0.0001. (**D**) Z-scored mean timecourse of -dt/t cGM, dt/t gPFI, and bsCSF from the same subject as in (**B**). (**E**) same as (**C**) for d/dt gPFI vs. bsCSF, t = 13.6, P < 0.0001. (**F**) Multi-level mediation analysis between d/dt cGM, d/dt gPFI and bsCSF shows a highly significant indirect path. ab = -3.64, P < 0.0001. (**G**) For cGM amplitude see Panel A. (**H**) For correlation coefficients d/dt GM-bsCSF see Panel A.
