## Supplementary figures and images for "Cerebrospinal fluid flow and clearance driven by lateral ventricle volume oscillations"

### Movie S1

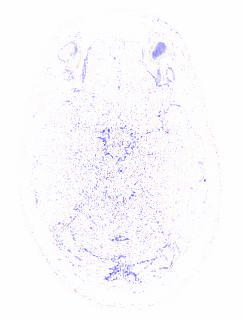

### Movie S2

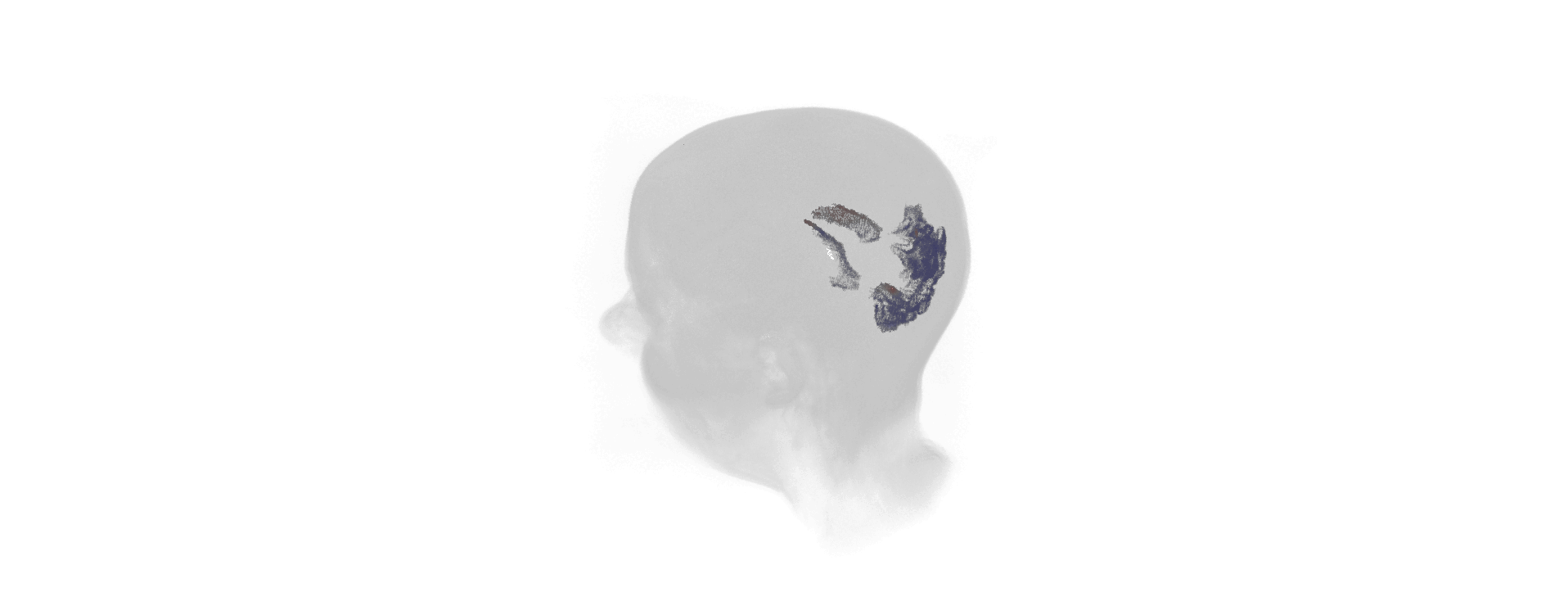
